# Synthetic algal-bacteria consortia for space-efficient microalgal growth in a simple hydrogel system

**DOI:** 10.1101/2021.03.01.433490

**Authors:** Noah Martin, Tatum Bernat, Julie Dinasquet, Andrea Stoftko, April Damon, Dimitri D. Deheyn, Farooq Azam, Jennifer E. Smith, Matthew P. Davey, Alison G. Smith, Silvia Vignolini, Daniel Wangpraseurt

## Abstract

Photosynthetic microalgae are an attractive source of food, fuel or nutraceuticals, but commercial production of microalgae is limited by low spatial efficiency. In the present study, we developed a simple photosynthetic hydrogel system that cultivates the green microalga, *Marinichlorella kaistiae* KAS603, together with a novel strain of the bacteria *Erythrobacter* sp.. We tested the performance of the co-culture in the hydrogel using a combination of chlorophyll-*a* fluorimetry, microsensing and bio-optical measurements. Our results showed that growth rates in algal-bacterial hydrogels were about 3-fold enhanced compared to hydrogels with algae alone. Chlorophyll-*a* fluorimetry based light curves found that electron transport rates were enhanced about 20% for algal-bacterial hydrogels compared to algal hydrogels for intermediate irradiance levels. We also show that the living hydrogel is stable under different environmental conditions and when exposed to natural seawater. Our study provides a potential bio-inspired solution for problems that limit the space-efficient cultivation of microalgae for biotechnological applications.

## 1. Introduction

Microscopic photosynthesizing algae produce a range of high value products including lipids and pigments (Borrowitzka 2013). Additionally, algal biomass is of great interest for use as feedstocks in aquaculture and for the generation of biofuels (Villarruel-Lopez et al. 2017, Khan et al. 2018). However, commercial large-scale production of microalgae is still limited by low spatial efficiency and associated high production and processing costs (e.g. Borrowitzka and Vonhak 2017). Algal cultivation techniques can generally be divided into open pond systems, closed photobioreactors and biofilm-based systems (Posten 2009). Open pond systems cultivate algae in raceway ponds and have low maintenance cost but generate only limited biomass per area (Tan et al. 2020). Photobioreactor systems allow for controlled conditions of irradiance, gas flux and temperature, and yield higher algal growth efficiencies, but have high operation and maintenance costs (Tan et al. 2020; Lee 2001). Biofilm-based systems cultivate algae as surface-attached biofilms rather than in liquid suspensions. Algal biofilm cultivation can lead to reduced operation costs due to limited water and energy use, as well as improved algal harvesting efficiencies (Ozkan et al. 2012; Berner et al. 2015). Biofilm systems also demonstrate greater CO_2_ utilization efficiency, and reduced harvesting cost (Blanken et al. 2016; Roostaei et al. 2018). These systems, however, are also constrained, often relying on sophisticated artificial architectures to compete with the efficiency of natural systems and are much harder to scale-up.

More recently, algae have also been cultivated while immobilized in hydrogels (Berner et al. 2015). Hydrogel immobilization enables reduced water usage during algal cultivation and provides a potential physical barrier against bacterial infections (Brenner et al. 2008; Covarrubias et al. 2012; He et al. 2016). 3D bioprinting has been used to create different hydrogel structures growing a range of microalgal strains (Krujatz et al. 2014; Lode et al. 2015; Wangpraseurt et al. 2020). To optimize light propagation in hydrogels with high microalgal densities, coral-inspired biomaterials have recently been developed (Wangpraseurt et al. 2020). However, the cultivation of microalgae in hydrogel-based systems, still requires further development regarding the exchange of gases and metabolites that are essential for microalgal growth (Podola et al. 2017).

To overcome diffusion limitation in attached cultivation systems, previous efforts have included the development of porous substrate-based bioreactors that make use of a porous membrane to deliver nutrients and promote gas exchange, whilst the surface of the biofilm is in direct contact with the ambient gas phase (Podola et al. 2017). In nature, benthic photosynthetic animals have faced similar challenges and photosynthesis in thick coral tissues could theoretically become limited by the diffusion-limited provision of HCO_3_^-^ from the ambient water phase. However, it has been shown that coral animal and bacterial respiration promote photosynthesis of their symbiotic microalgae, suggesting that the coral host provides essential metabolites and nutrients locally to the microalgae (e.g. Kuhl et al. 1996; Schrameyer et al. 2014).

In corals, the microbial community performs critical functions for the coral holobiont including pathogen protection, sulfur and nitrogen cycling as well as beneficial modulations of the host microhabitat (Rosenberg et al. 2009; Krediet et al. 2013; Ceh et al. 2013). Benefits of bacterial communities for an algal host have been documented in free-living algae as well (e.g. Kazamia et al. 2012). Some bacteria can provide a local supply of essential nutrient compounds required by the algae, including nitrogen, inorganic carbon, vitamin B12 (cobalamin), and growth promoting hormones (Kouzuma and Watanabe 2015). For example, one study estimated that 50% of algal species are cobalamin auxotrophs, implying a reliance on bacterial-produced cobalamin (Croft et al. 2005). More generally, symbiotic relationships between microalgae and bacteria often employ a mutually beneficial exchange of carbon and nitrogen (Thompson et al. 2012). Experiments working with the microalgae *Chlorella* in co-culture with a known growth promoting bacteria in alginate beads demonstrates enhanced growth which can be utilized for biotechnological applications (Gonzalez and Bashan 2000). Accordingly, there is a growing interest in using microbial consortia for enhanced biomanufacturing (Padmaperuma et al. 2018; Nai and Meyer 2018).

Here, we developed a novel gelatin-based hydrogel system by combining microalgae and bacteria for space-efficient microalgal cultivation. We chose the green microalga *Marinichlorella kaistiae* KAS603 and screened 14 marine bacterial strains for beneficial effects on algal biomass. Based on these results, we further measured the bio-optical properties and photosynthetic performance of a synthetic co-culture between *Marinichlorella kaistiae* KAS603 and a novel strain of *Erythrobacter* sp.. We then evaluated the beneficial effects of the *Erythrobacter* strain on a range of microalgae covering coccolithophorids, red algae and other species of green microalgae. Finally, the mechanical stability of our hydrogel system was tested under different environmental conditions.

## 2. Methods

### Experimental approach

To test for beneficial effects of algal-bacterial co-culture, we assessed a range of bacterial and algal strains. *Marinichlorella kaistiae* KAS603 (Sánchez-Alvarez et al. 2017) was used as model algal strain. *M. kaistiae* KAS603 is a robust algal strain that is morphologically similar to *Chlorella* and has high lipid and biomass production rates (Sánchez-Alvarez et al. 2017). *M. kaistiae* KAS603 has been successfully grown in 3D bioprinted gelatin-based hydrogels (Wangpraseurt et al. 2020). The beneficial impact of 14 different bacterial strains (see Table S1) on *M. kaistiae* KAS603 growth was investigated over 3-day co-culture experiments. These preliminary experiments suggested enhanced growth with the strain SIO_La6, closely related to *Erythrobacter* sp., (Table S1), which was then used as our bacterial model for co-culture experiments. Finally, to test whether these beneficial effects of SIO_La6 were transferrable to other microalgal species, co-cultures between SIO_La6 and *Micromonas sp., Porphyridium cruentum, Pleurochrysis carterae,* and *Amphidinium carterae,* were also investigated. Co-culture experiments with *M. Kaistiae KAS603* were conducted also in liquid culture to assess the relative effect of algae immobilization in hydrogels (Fig. S1).

### Stock cultures

Bacterial stock cultures were cultivated in Zobell broth at 25°C under sterile conditions. Bacterial cultures used for hydrogel immobilization were harvested during exponential growth in Zobell broth as determined via optical density (OD) measurements (Begot et al. 1996) and flow cytometry (Gasol and Del Giorgio 2000). Bacterial cultures were identified by16S rDNA Sanger sequencing (using the primer pair 27F-1492R) to determine their closest phylogenetic relations (Table S1). Algal stock cultures were grown in artificial seawater medium (ASW, Darley and Volcani 1969) at 25 °C under a continuous irradiance regime of 150 μmol photons m^-2^ s^-1^ provided by white LED light panels (AL-H36DS, Ray2, Finnex). Microalgae were harvested from liquid stock cultures in the exponential growth phase for hydrogel immobilization. Cell density was measured using a hemocytometer, with 3 technical replicate counts per algal stock sample.

### Algal-bacterial hydrogel fabrication and cultivation

Hydrogels were made by using a 10% solution of porcine gelatin (type-A, Sigma-Aldrich, USA) in ASW. The solution was prepared by heating the gelatin-ASW mixture on a hot plate under continuous stirring to 90 °C until it was optically clear. The solution was cooled to 30 °C and 2.5 mL of the gel solution was rapidly mixed with 2 mL of the algal stock solution (at a concentration of 1.36 x 10^7^ cells/mL for main *M. kaistiae* experiments) and 0.5 mL of either sterile Zobell medium (for monoculture control gels) or Zobell medium containing a chosen bacterial strain (for co-culture gels) (Figure 1). Bacterial density for cultivation experiments was chosen at an OD600 of 0.02. We also performed preliminary growth experiments using different starting concentrations of microalgal cell density (Fig. S2). The solution was vortexed for 30 s, to ensure proper mixing of algae and bacteria, before it was poured into Petri dishes. Gelation was facilitated by keeping the Petri dishes at 18 °C for 1 hour, which resulted in gels that were ~10 mm thick. Gels were then cultivated at 25 °C under a continuous irradiance regime of 150 μmol photons m^-2^ s^-1^ provided by white LED light panels (AL-H36DS, Ray2, Finnex).

**Fig. 1.**
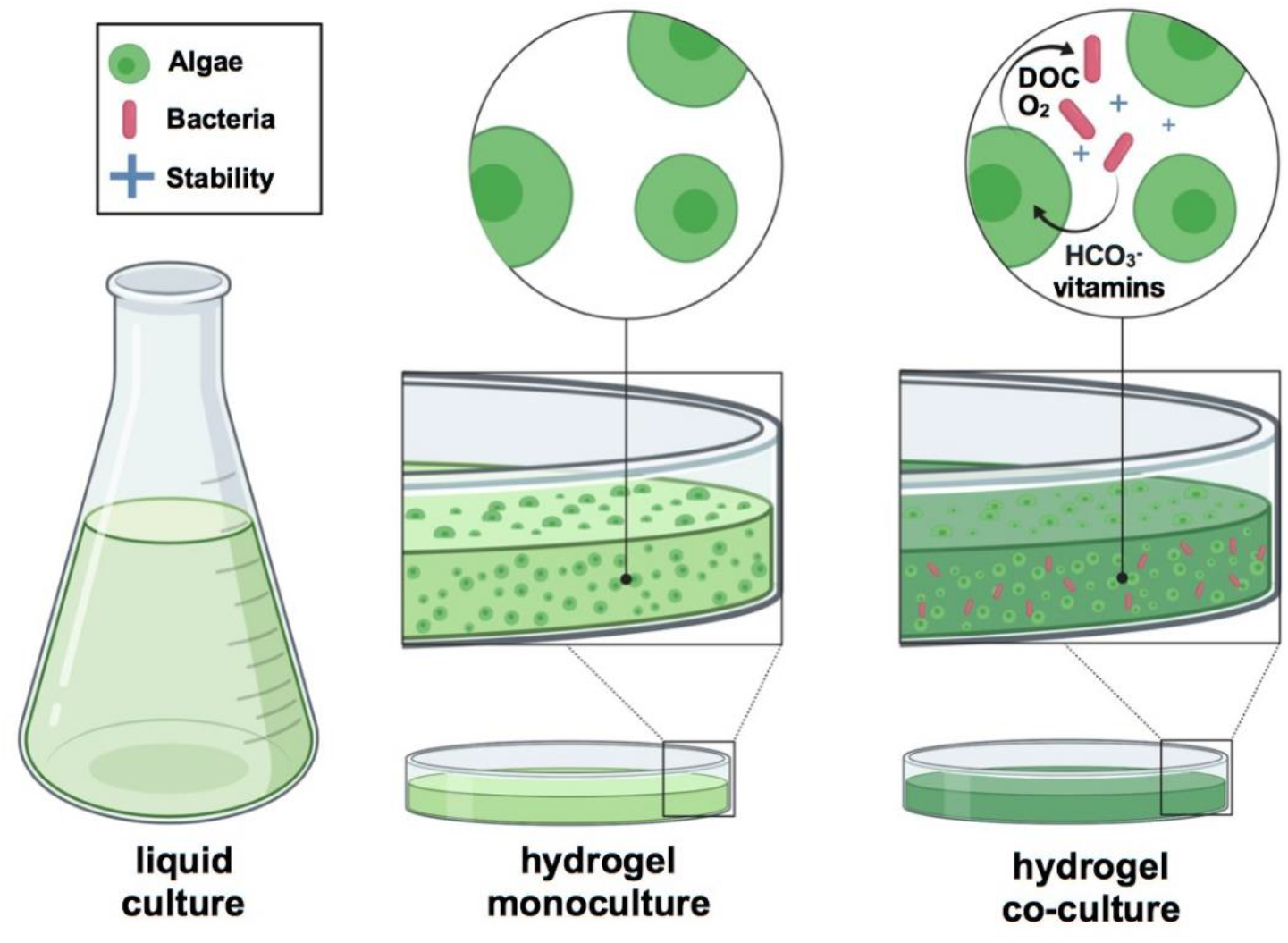
Development of a synthetic co-culture between microalgae and *Erythrobacter sp.* in a gelatin-based hydrogel. Algae were grown in monoculture and in co-culture with *Erythrobacter sp.* both in liquid culture and in hydrogel configuration. Arrows indicate potential interactions between algae and bacteria that were hypothesized to enhance algal growth. Microalgal photosynthesis generates O_2_ and dissolved organic carbon (DOC) that fuels bacterial metabolism. In turn, bacterial activity provides an inorganic carbon source for photosynthesis and vitamins. This synthetic co-culture enhances the stability of the biopolymer when exposed to potential pathogens. (Figure was created with BioRender.com)

### Performance testing

#### Microalgal cell density

Hydrogels were liquified by heating to 30°C on a hot plate. The liquid algal suspension was then diluted with ASW and the cell density was determined with a hemocytometer (see above). The accuracy of this approach was tested using stock cultures of known cell density, showing an error of less than 3% between expected and measured cell densities.

#### O_2_ microsensor measurements

Clark-type O_2_ microsensors (tip size of 25 μm, a 90% response time of <0.5 s and a stirring sensitivity of ~1%; Unisense A/S, Aarhus, Denmark) were used to measure O_2_ production and consumption of the hydrogels as described previously (Wangpraseurt et al. 2012). Briefly, microsensors were connected to a picoammeter (Unisense, Denmark) and operated by an automatic microsensor profiler (MU1, Pyroscience GmbH, Germany). Hydrogels were placed in a black acrylic flow chamber and flowing seawater was supplied at a flow velocity of 0.5 cm s^-1^ at 25° C and a salinity of 35. Microsensors were positioned at the surface of the hydrogel by observing the microsensor tip with the aid of a dissecting microscope and the use of an automated micromanipulator (MU1, Pyroscience GmbH, Germany). Steady-state O_2_ concentration profiles from the hydrogel surface through the diffusive boundary layer (DBL) and into the mixed turbulent water phase above were performed in 100 μm steps under an incident photon irradiance of Ed(PAR) = 0 and 550 μmol photons m^-2^ s^-1^. O_2_ microsensors were linearly calibrated from readings at 100 % air saturated seawater at experimental temperature and using anoxic water (flushed with N2). Percent air saturation in seawater at experimental temperature and salinity was transformed to O_2_ concentration (μmol O_2_ L^-1^) using gas tables (Ramsing and Gundersen 2011).

#### Variable chlorophyll a fluorimetry

We used a variable chlorophyll a fluorometer (diving PAM II, Walz, Germany) to characterize PS II performance (Baker 2008). The fiber of the PAM system was mounted on a laboratory stand and directed vertically towards the surface of the hydrogels at a fixed distance of 1 cm. Hydrogels were dark adapted for at least 30 minutes before experimental measurements. Rapid light curves (RLC) (Ralph and Gademan 2005) were performed over a range of 8 irradiance intensities spanning 0-1500 umol photons m^-2^ s^-1^ of incident downwelling irradiance. For each RLC, the dark-adapted hydrogels were incubated at each experimental irradiance regimes for 15 seconds followed by a saturation pulse. The variable chlorophyll fluorescence data was analyzed as described previously (Ralph and Gademan 2005). Briefly, the maximum quantum yield of PSII was calculated as:

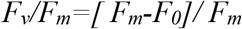

and the effective quantum yield of PSII was calculated as:

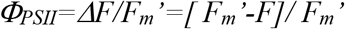

Where F_0_ and F describe the minimum and transient fluorescence and F_m_ F_m_’ describe the maximum fluorescence in the light adapted state. The electron transport rate was calculated as ETR= Φ_PSII_ x E_d_ X 0.5 x AF, where E_d_ is the incident downwelling irradiance (400–700 nm), 0.5 assumes the equal distribution between PSI and PSII and AF denotes the absorption factor which was assumed to be 0.83 (Ralph and Gademan 2005). It is important to note that AF will vary as a function of pigment and cell density and thus serves only as an approximation (Wangpraseurt et al. 2019). The photosynthetic light curves were fitted to the empirical equations of Platt and Gallegos (1980), using a Marquardt-Levenberg regression algorithm:

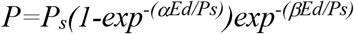

where *Ps* is a scaling factor defined as the maximum potential rETR, *α* describes the light use efficiency, i.e. the initial slope of the RLC and *β* characterizes photoinhibition and indicates the slope of the RLC where PSII declines. The maximum electron transport rate ETR_max_ and the light intensity at half saturation, Ek were calculated as:

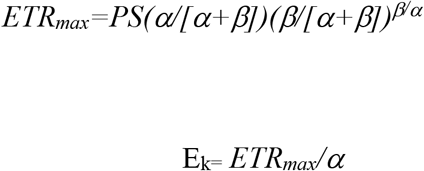

The fitting procedure was sensitive to initial guesses of PS, *α, β,* which were adjusted for each curve fitting. All fitting was done with custom codes written in Matlab 2018b.

#### Bio-optical properties of the hydrogels

Irradiance reflectance of the gels were measured with a 0.7 mm wide flat-cut fiber-optic reflectance probe (Ocean Optics, USA) with the hydrogels positioned in the black acrylic flow-through system described previously. The hydrogel was illuminated vertically incident by a light source emitting broadband white light. Reflectivity was determined with the reflectance probe positioned at a distance of 500 μm from the hydrogel surface. All reflectivity measurements were normalized to the reflectivity of a 10%, 20% and 99% white diffusing reflectance standard (Spectralon, Labsphere, USA). These measurements occurred under identical configuration and distance to light source as on the hydrogel surface, but were performed in air. Measurements of scalar irradiance (i.e. the integral quantum flux from all directions around a given point) were measured with fiber-optic microsensors (zensor, Denmark) as described previously (Wangpraseurt et al. 2012).

### Bacterial contamination experiment

To test whether the co-culture with *Erythrobacter* sp. SIO_La6 strain would provide protection from other microbes, we exposed the hydrogels to natural unsterilized seawater supplied from the Scripps Pier. For these tests, 3-day old hydrogels were incubated with the natural seawater for 1.5 hours in a beaker under low turbulent flow. The gels were then removed, and cultivation in the environmental growth room continued as described above. The gels were visually examined at every day after exposure and photographed to assess visual differences, such as noticeable cell death, bacterial growth or hydrogel liquification, indicative of gelatin-degrading bacteria.

## Results and Discussion

Here, we developed a simple hydrogel system for the space-efficient co-culture of microalgae. We found that a novel strain of *Erythrobacter* sp. (SIO_La6, Fig. 2) isolated from Southern California coastal waters (off Scripps Pier) has beneficial effects on growth and photosynthetic performance of microalgae immobilized in hydrogels.

**Fig. 2.**
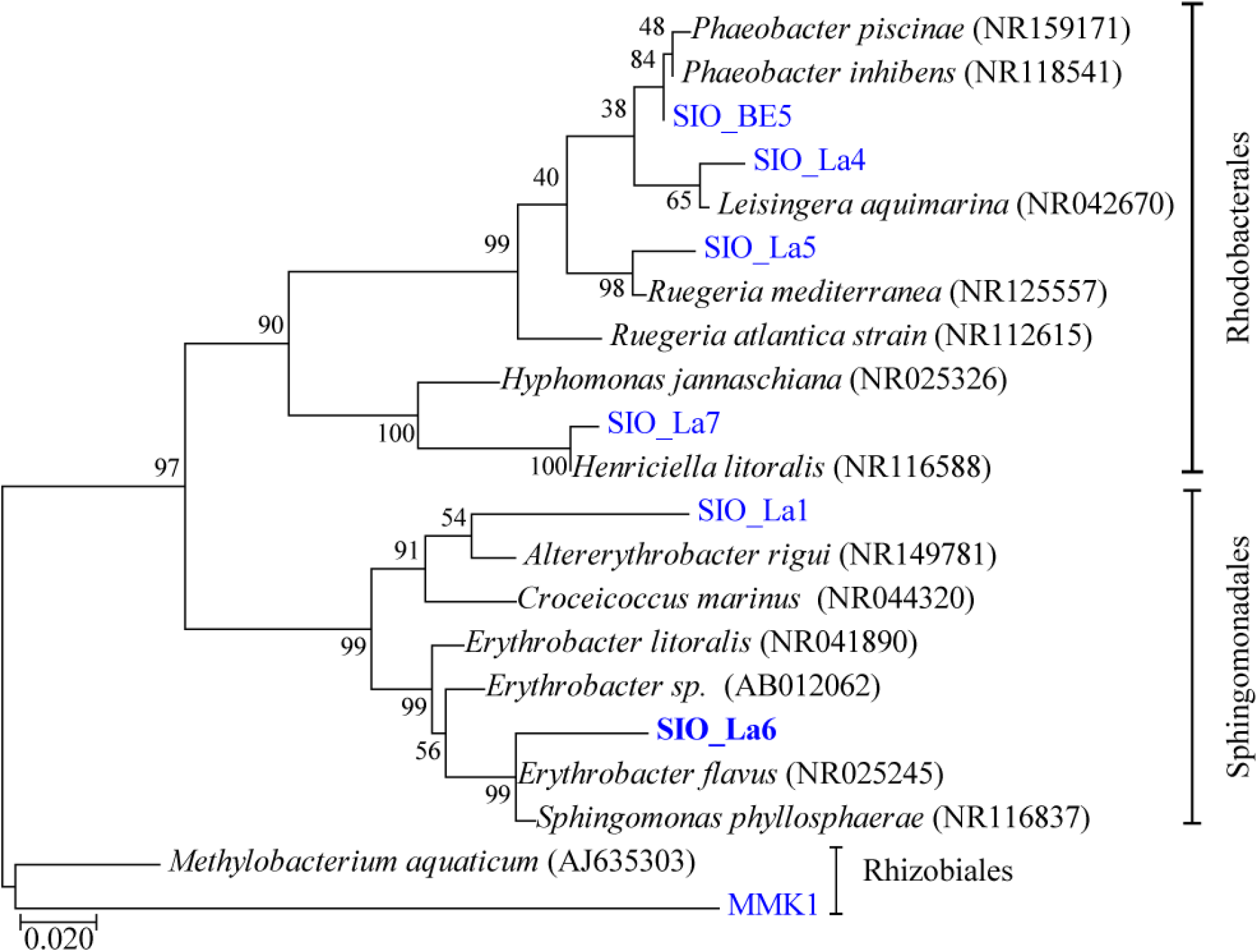
Maximum likelihood tree of Alpha-proteobacteria sequences closely related to the tested isolates (SIO_La6). Reference sequences from NCBI are indicated in italic. Bootstrap values (n=1000) are indicated at nodes; scale bar represents changes per position.

### Cell density differences between treatments

Microalgal cell density was on average 2.3-fold enhanced for *Marinichlorella kaistiae* KAS603 gels co-cultured with SIO_La6 (mean = 2.85 x 10^7^ cells mL^-1^, SD = 5.94 x 10^6^, *n* = 5) compared to monoculture gels (1.18 x 10^7^ cells mL^-1^, SD = 4.06 x 10^6^, *n* = 5) after 72 h of cultivation (paired t-test, p <0.001, Fig. 3A). The cell doubling time was 16.75 h for co-cultures compared to 33.11 h for monocultures (Fig. 3). The beneficial effects of co-culture with *Erythrobacter sp.* SIO_La6 were also evident in liquid culture, although the relative growth stimulating effect was 15% higher in hydrogel (Supplementary Fig. 2). In a stagnant hydrogel, gas exchange is likely to become a limiting growth factor, while such limitation is unlikely to occur in a liquid mixed culture. Thus, the relative enhancement for hydrogel cultures could suggest that bacterial colonies stimulate gas exchange, and provide nutrients locally within the hydrogel. Indeed, bacteria observed during confocal microscopy were observed forming aggregates around algal cells (Supplementary Fig. 3). This proximity, and reduced diffusion in a gel compared to liquid culture, may account for better access of algae to growth enhancing nutrients from bacteria in co-cultured hydrogels.

**Fig. 3.**
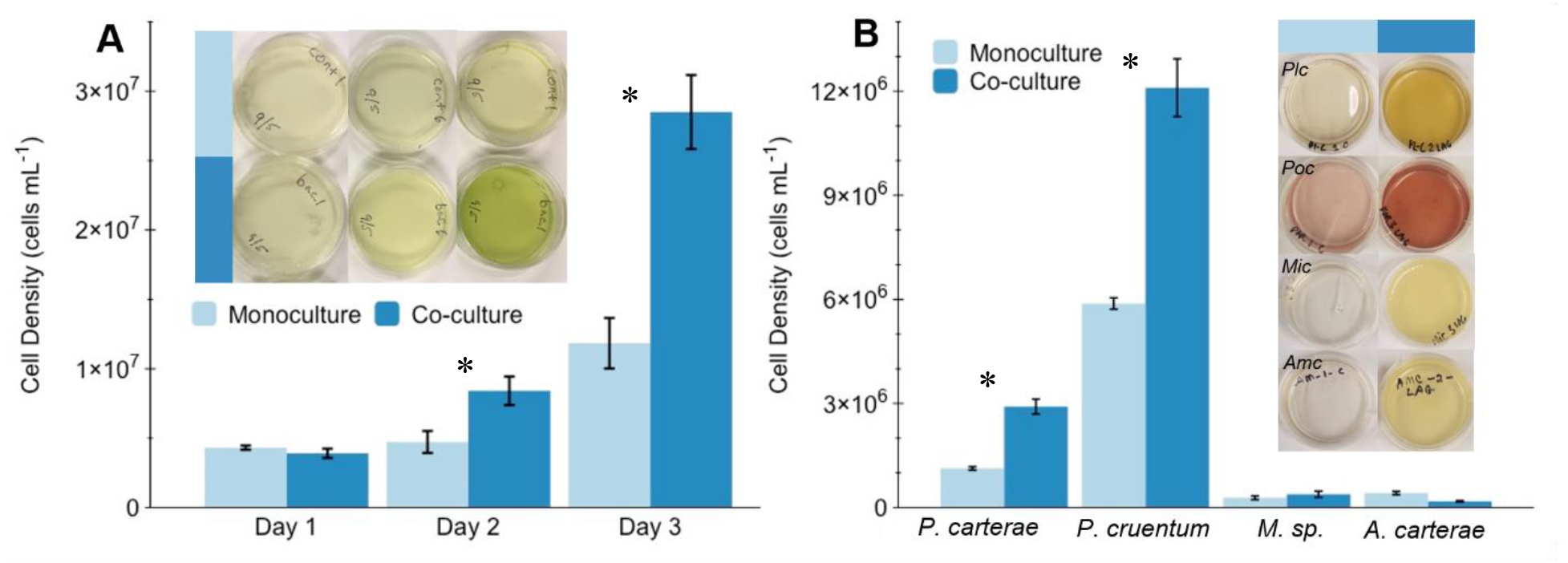
Effect of algal-bacterial hydrogel co-culture on microalgal cell density growth. (**A**) 3-d growth dynamics of *Marinichlorella kaistiae* KAS603 in monoculture (light blue) and in co-culture with *Erythrobacter* sp. SIO_La6 (dark blue). Insets show example top view images of hydrogels each day. Data are means ±SD, *n* = 15. (**B**) Cell density of *Pleurochrysis carterae, Porphyridium cruentum, Micromonas sp. and Amphidinium carterae* after 3 days of growth in monoculture and co-culture. Images show top view images of hydrogel after 3 days. Data are means ± SD *n*=2. * indicates a significant difference between treatments (p<0.05, paired student’s t-test).

Following the successful tests with *M. kaistiae* KAS603, other common microalgae were tested in co-culture with SIO_La6. The bacterial co-culture enhanced microalgal growth for three of the five microalgal strains compared to monoculture controls (Fig. 3B). Cell densities after 3-d of cultivation were at least 2-fold higher for the coccolithophorid algae *Pleurochrysis carterae* and the red algae *Porphyridium cruentum* when grown in co-culture hydrogels (Fig. 3B). Interestingly, cultures that did not perform well in co-culture (e.g. *Micromonas sp.* and *Amphidinium carterae*) also showed limited growth when encapsulated in the gelatin-based hydrogel in monoculture, suggesting that hydrogel immobilization interfered with the growth dynamics of these algae (Fig. 3B). This suggests that *Micromonas sp.* and *Amphidinium carterae* might not be suitable candidates for biotechnological applications using hydrogel immobilization.

### Co-culture effects on microalgal photosynthesis and bio-optics

Compared to *Marinichlorella kaistiae* KAS603 monocultures, O_2_ microsensor measurements in co-cultures indicated 4.9-fold enhancements of net photosynthesis at high light (550 umol photons m^-2^ s^-1^) irradiance regimes (Fig. 4A). Additionally, co-cultures exhibited about 4.3-fold greater rates of dark respiration (Fig. 4A). Variable chlorophyll-*a* fluorimetry measurements showed significant enhancements in the maximum quantum yield of PSII (Fv/Fm) for co-culture hydrogels compared to monoculture hydrogels during 7 days of growth (mean = 0.603, SD = 0.022 vs mean = 0.535, SD = 0.004, respectively; Fig. 4B, paired t-test p = 0.006). Fv/Fm is a key parameter used to assess the healthiness of photosynthesizing microalgae (e.g. Baker 2008) and thus suggests that algae in co-culture displayed superior photosynthetic capacities. Likewise, relative electron transport rates showed clear differences in key photosynthetic parameters including α and ETRmax (Fig. 4D-F, Table 1). For instance, at day 3 ETRmax was about 71.6% higher for co-cultures vs monocultures Fig. 4D-F, Table 1).

**Fig. 4.**
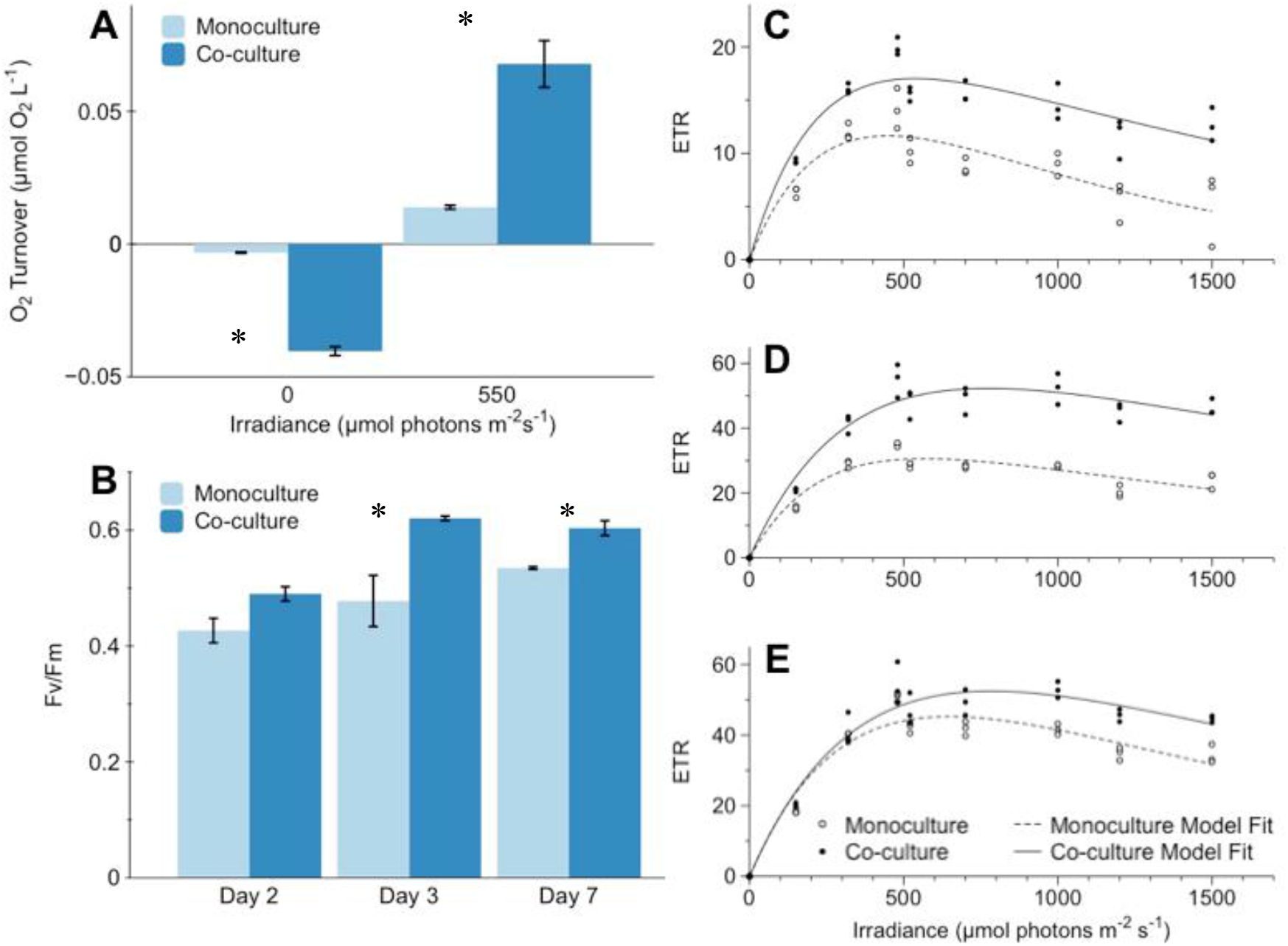
Photosynthetic performance of hydrogels in mono- and co-culture. (**A**), O_2_ turnover based on O_2_ microsensor measurements of the linear O_2_ flux from the surface into the diffusive boundary layer performed at 0 (dark respiration) and at 550 μmol photons m^-2^ s^-1^ (net photosynthesis. (**B**) Maximum quantum yield of PSII (Fv/Fm) and electron transport rates (ETR) at (**C**) day 2, (**D**) day 3, and (**E**) day 7 of algal cultivation. Data are means ± SD (*n* = 4 for panel A and *n*= 3 for panel B-E). Note that y-axis scale was adjusted for clarity in panel C-E. * indicates a significant difference between treatments (p<0.05, paired student’s t-test).

Although areal net photosynthetic (P_n_) rates were strongly enhanced in co-culture, these differences were also affected by the greater algal growth in co-culture (Fig. 3). However, normalizing P_n_ rates to the differences in biomass still suggests an approximate doubling in net photosynthesis in co-culture vs monoculture (compare Fig. 3A and 4A). As *Erythrobacter sp.* are anoxygenic phototrophic bacteria and thus does not produce O_2_ (Koblizek et al. 2003) such differences strongly suggest cell specific enhancements of photosynthetic activity by *M. kaistiae* KAS603 in the presence of *Erythrobacter*. It is important to note that these measurements include respiratory activity by the bacteria, further strengthening the argument of enhanced algal photosynthesis in co-culture. PAM measurements can detect potential electron transport by *Eyrythrobacter* sp. (Chandaravithoon et al. 2020), however we did not find any measurable quantum yield of PSII from SIO_LA6 in monoculture (F_v_/F_m_=0, data not shown). Additionally, diffuse reflectance measurements did not show characteristic absorption peaks of bacteriochlorophyll *a* at ~ 750 nm (Fig. 5, Yurkov and Beatty 1998), suggesting that pigment synthesis and photosynthetic electron transport might be low by this *Erythrobacter* strain. In turn, reflectance in the near-infrared region (~750 nm) was about 2.5-fold enhanced which could be indicative of the production of light scattering microbial extracellular polymeric substances (EPS, Flemming and Wingender 2001). Such EPS has previously been shown to scatter light and could potentially enhance the internal actinic irradiance intensity which would further promote photosynthesis (Decho et al. 2003; Fisher et al. 2019). Clearly, there are various potential mechanisms underlying the enhanced photosynthetic performance of the co-culture hydrogels and a detailed understanding of the mechanisms was beyond the scope of this first study. However, taken together our results indicate that *Erythrobacter* sp. SIO_La6 enhances *Marinichlorella kaistiae* KAS603 photosynthesis (Table 1) which could explain the enhanced algal biomass in co-culture.

**Fig. 5.**
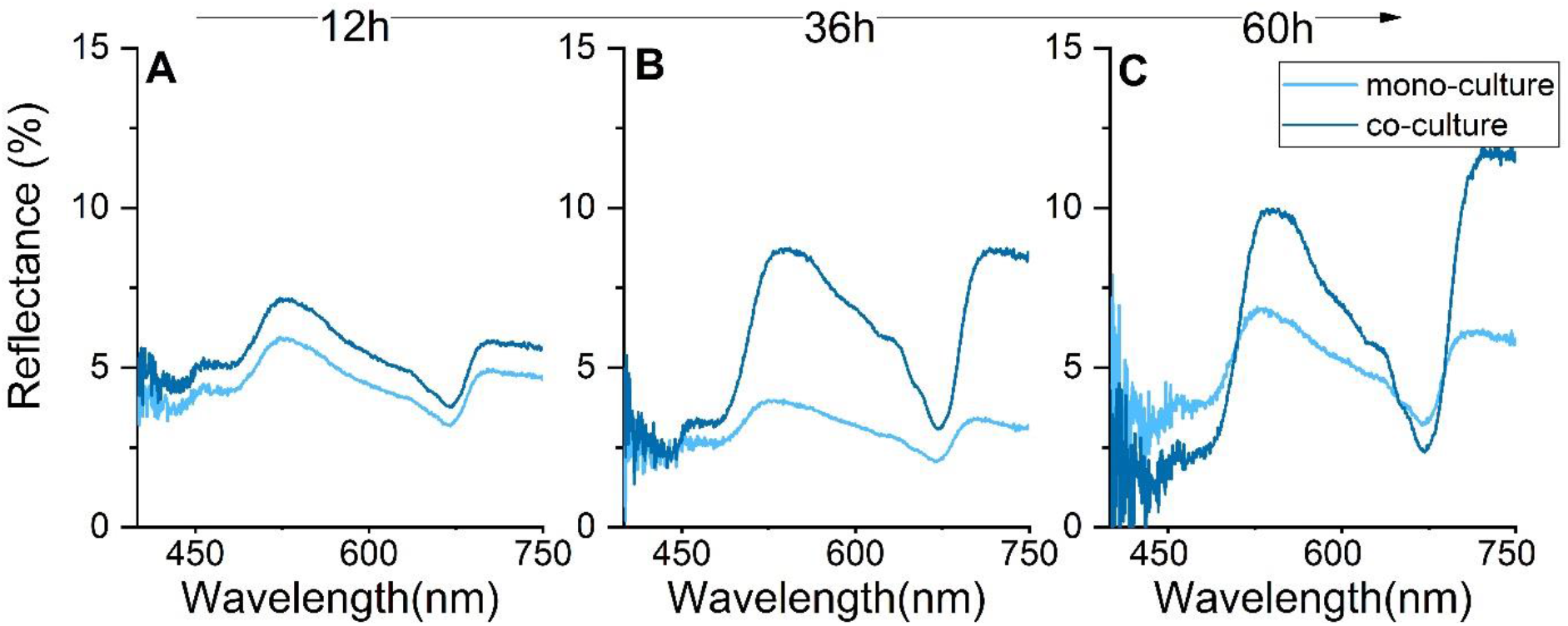
Hydrogel diffuse reflectance (%) after (A) day 1, (B) day 2, and (C) day 3 of algal cultivation. Data are means from 3 hydrogels, error bars are omitted for clarity (SD was less than 5%)

**Table 1.** Photosynthetic performance of *Marinichlorella kaistiae* KAS603 grown in the hydrogel alone (mono-culture) or together with *Erythrobacter* sp. SIO_La6 (co-culture). Parameters are derived from the best fit from all replicate measurements (*n*=3, lines in Fig. 4 C-E)

|  | Day 2 |  | Day3 |  | Day7 |  |
| --- | --- | --- | --- | --- | --- | --- |
|  | Mono-culture | co-culture | Mono-culture | co-culture | Mono-culture | co-culture |
| $\alpha$ | 0.07 | 0.10 | 0.17 | 0.21 | 0.21 | 0.20 |
| $\beta$ | 0.04 | 0.015 | 0.03 | 0.034 | 0.06 | 0.05 |
| $ETR_{max}$ | 11.64 | 17.03 | 30.59 | 52.30 | 45.26 | 52.5 |
| $E_k$ | 158 | 169 | 180 | 245 | 220 | 261 |
| $R^2$ | 0.8 | 0.90 | 0.91 | 0.93 | 0.93 | 0.94 |

### Contamination resistance in hydrogels

A potential key problem in cultivating microalgae in hydrogels is that most biopolymers are readily degraded by various bacterial communities (Pathak et al. 2017). We hypothesized that co-cultivation might provide protection from such degradation by occupying microbial habitats within the hydrogel and potentially producing antibiotics. Such concept is analogous to the role of the microbial community in the coral mucus, which protects from opportunistic microbes (Shnit-Orland et al. 2009). Following exposure to natural seawater, co-culture gels remained viable and no visible degradation of the gelatin matrix was noticeable even after 7 days of cultivation (Fig. 6A-E). However, monocultures showed clear degradation and liquefaction of the polymer matrix within 24 hours (Fig. 6A-E). Likewise, previous experiments using *Chlorella*-bacteria co-cultures in alginate beads found reduced contamination by foreign bacteria from the environment and concluded that co-cultured bacteria provide a physical barrier (Covarrubias et al. 2012). Here, it is likely that DOC produced by the algae might enhance virulence factors (present in SIO_La6 genomes, J. Dinasquet pers. com.) and toxin production as observed in other Erythrobacter species in the presence of algal DOC (Cardenas et al. 2018). This induced pathogenicity might have antagonistic effects against environmental contaminants. Although the mechanisms warrant further investigation, these initial results suggest protective effects of our synthetic co-culture hydrogel from external microbes. This could be further developed as a viable bio-inspired alternative to costly antibiotic treatments that are currently used in such cultivation approaches (Berner et al. 2015).

**Fig. 6.**
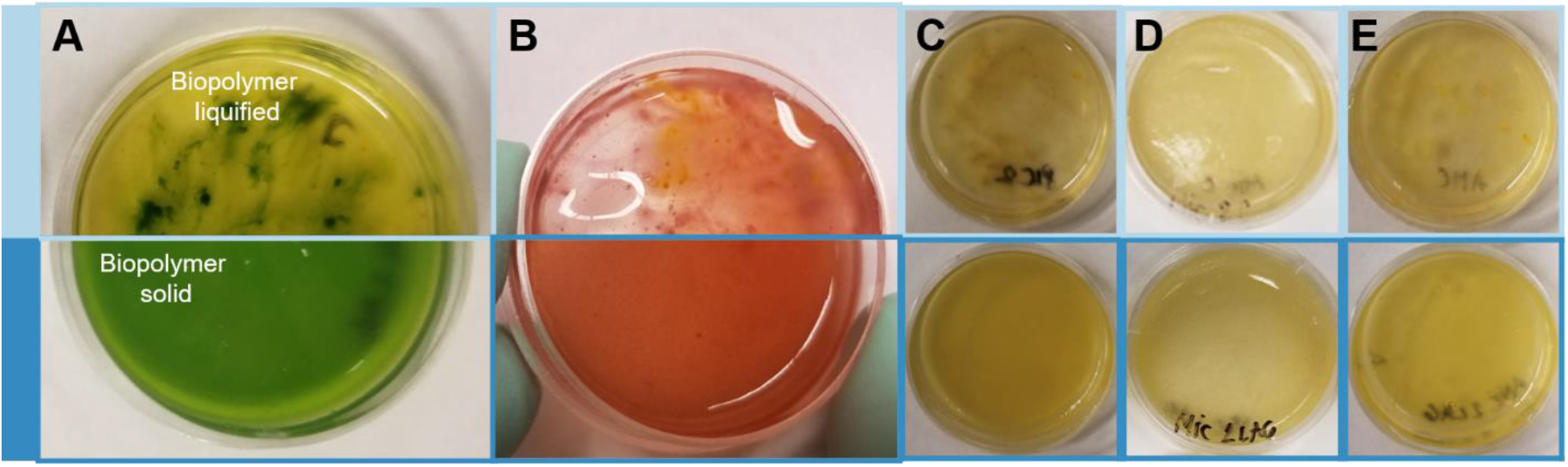
Biopolymer stability after exposure to natural seawater. Images show top view of hydrogels after 7 days of the seawater exposure experiment. Monocultures (top panels, light blue) are liquified while co-cultures remain solid (bottom panels, dark blue) for (A) *Marinichlorella kaistiae* KAS603 (**B**) *Porphyridium cruentum,* (**C**) *Pleurochrysis carterae.,* (D) *Micromonas sp.* and (E) *Amphidinium carterae*

## Conclusions

This study developed a simple hydrogel system for microalgal cultivation in co-culture with a novel strain of *Erythrobacter* sp. Our findings demonstrate enhanced photosynthetic activity and growth rates of microalgae in co-culture when immobilized in our hydrogel system. We further show that our gelatin-based hydrogel is easy to fabricate, requires low maintenance, and remains stable when the co-culture is exposed to natural contaminants. Our study suggests that co-cultivation in hydrogels of microalgae with *Erythrobacter sp.,* enhances microalgal growth and density, and could potentially reduce the need for costly antibiotics. We conclude that hydrogel algal-bacterial co-culture is a simple, bio-inspired approach that can be further developed to solve some problems that currently limit microalgal cultivation.

## Data availability

All raw data generated during this study are deposited on figshare (link: XX).

## Author contributions

Conceptualized and designed the study: D.W., J.D., N.M., T.B., S.V., A.G.S., M.D., Performed experimental measurements: N.M., T.B., A.S., A.D. Analyzed and interpreted data: N.M., T.B. D.W. Provided reagents, materials and analysis tools: D.W., J.D., D.D.D., S.V., F.A., J.E.S., Supervised the study: D.W., S.V., D.D.D. Wrote the manuscript: N.M, T.B., DW. All authors critically assessed the results and edited drafts of the manuscript.

## Acknowledgements

Mark Hildebrand is thanked for initial input on the experimental approach and for providing *Marinchlorella* cultures. Orna Cook is thanked for providing algal cultures. Funding: This study was funded by the European Union’s Horizon 2020 research and innovation programme (702911-BioMIC-FUEL, D.W., D.D.D., S.V.), the European Research Council (ERC-2014-STG H2020 639088, S.V.), the European Union’s 7^th^ Framework programme (PIOF-GA-2013-629378, J.D.), the Gordon and Betty Moore Foundation (Grant GBMF4827,F.A.).

## Notes

### Competing Interest Statement

The authors have declared no competing interest.

