## Supplementary Information for "Synthetic algal-bacteria consortia for space-efficient microalgal growth in a simple hydrogel system"

*shared first author

#correspondance: (D. Wangpraseurt)

**Supplementary Table 1**

Bacterial isolates tested from our culture collection. Closest relative assignment based on 16S rRNA gene sequencing. All bacteria were isolated off Scripps Pier.

| **Bacterial strain** | **Closest relative** |
|---|---|
| SIO_BE5 | AlphaProteobacteria; Rhodobacterales; Rhodobacteraceae; *Phaeobacter* sp. |
| SIO_La7 | AlphaProteobacteria; Hyphomonodales; Hyphomonodaceae; *Henriciella* sp |
| SIO_La6 | AlphaProteobacteria; Sphingomonadales; Erythrobacteraceae; *Erythrobacter* sp. |
| SIO_La5 | AlphaProteobacteria; Rhodobacterales; Rhodobacteraceae; *Phaeobacter* sp. |
| SIO_La4 | AlphaProteobacteria; Rhodobacterales; Rhodobacteraceae; *Phaeobacter* sp. |
| SIO_La1 | AlphaProteobacteria; Sphingomonadales; Erythrobacteraceae; *Erythrobacter* sp. |
| B6P1 | Gammaproteobacteria; Alteromonadales; Alteromonadaceae; *Alteromonas macleodii* |
| AltSIO | Gammaproteobacteria; Alteromonadales; Alteromonadaceae; *Alteromonas macleodii* |
| P1RIIB2 | Bacteroidetes; Flavobacteriales; Flavobacteriaceae; *Cellulophaga* sp. |
| BBFL7 | Bacteroidetes; Flavobacteriales; Flavobacteriaceae; *Flavobacterium* sp. |
| DMS2 | Gammaproteobacteria; Oceanospirillales; Oceanospirillaceae; *Marinomonas* sp. |
| A1.2 | Bacteroidetes; Flavobacteriales; Flavobacteriaceae; *Polaribacter* sp. |
| MMK1 | Alphaproteobacteria; Rhizobiales; Methylobacteriaceae; *Methylobacterium* sp. |
| TW7 | Gammaproteobacteria; Alteromonadales; Pseudoalteromonadaceae; *Pseudoalteromonas* sp. |

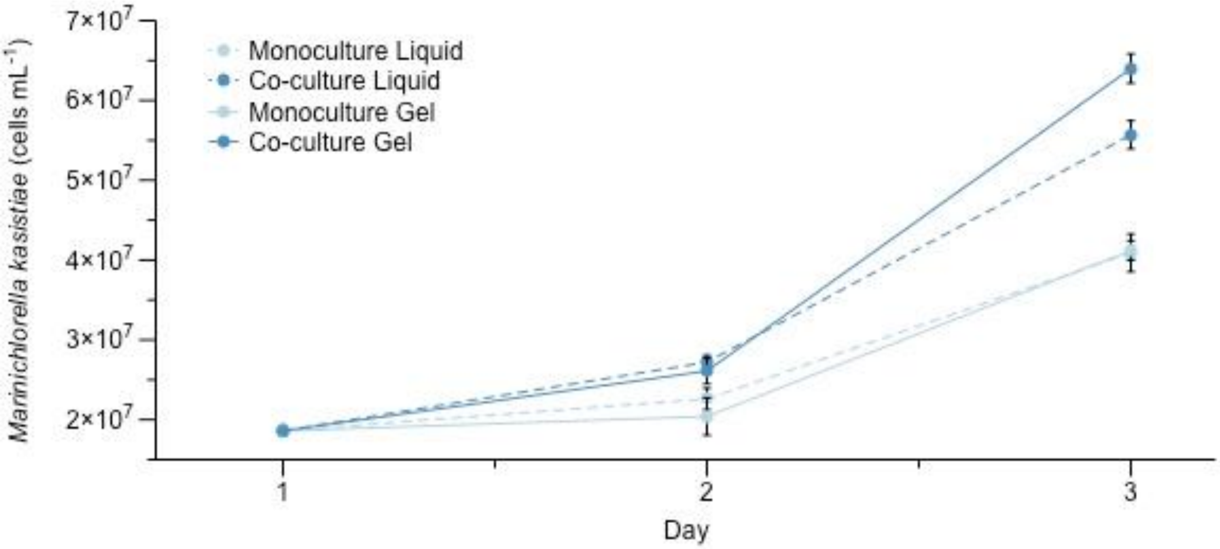


**Supplementary Fig. 1** Comparison of cell growth of *Marinichlorella kaistia*e KAS603 in co-culture with Erythrobacter strain SIO_LA6 (dark blue) and in mono-culture (light blue) when cultivated in liquid medium vs hydrogel. Data are means ± SE (*n* = 2-3 hydrogels/liquid culture flasks)

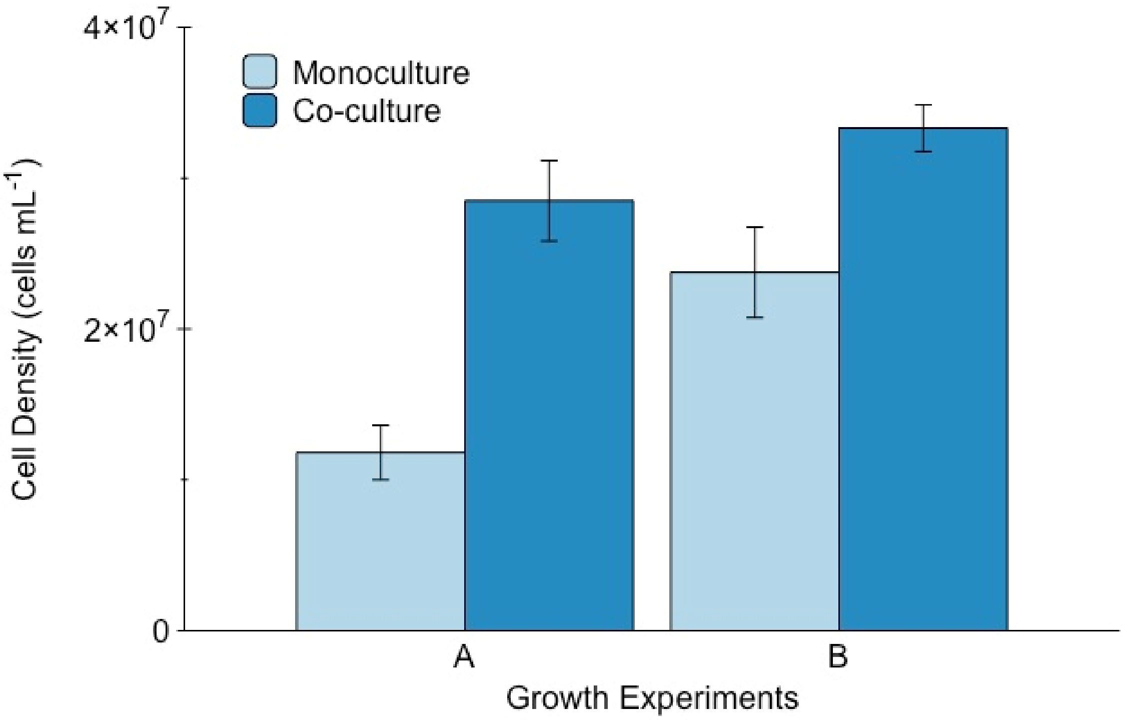


**Supplementary Fig. 2** Effect of algal-bacterial hydrogel co-culture on microalgal cell density growth after 72 hours. Microalgal starting cell density was 5.42 x $10^6$ (± 4.2 x $10^5$) cells/mL for growth experiment A and 2.21 x $10^7$ (± 4.9 x $10^5$) cells/mL for growth experiment B. Data are means ± SE (*n* = 3 hydrogels).

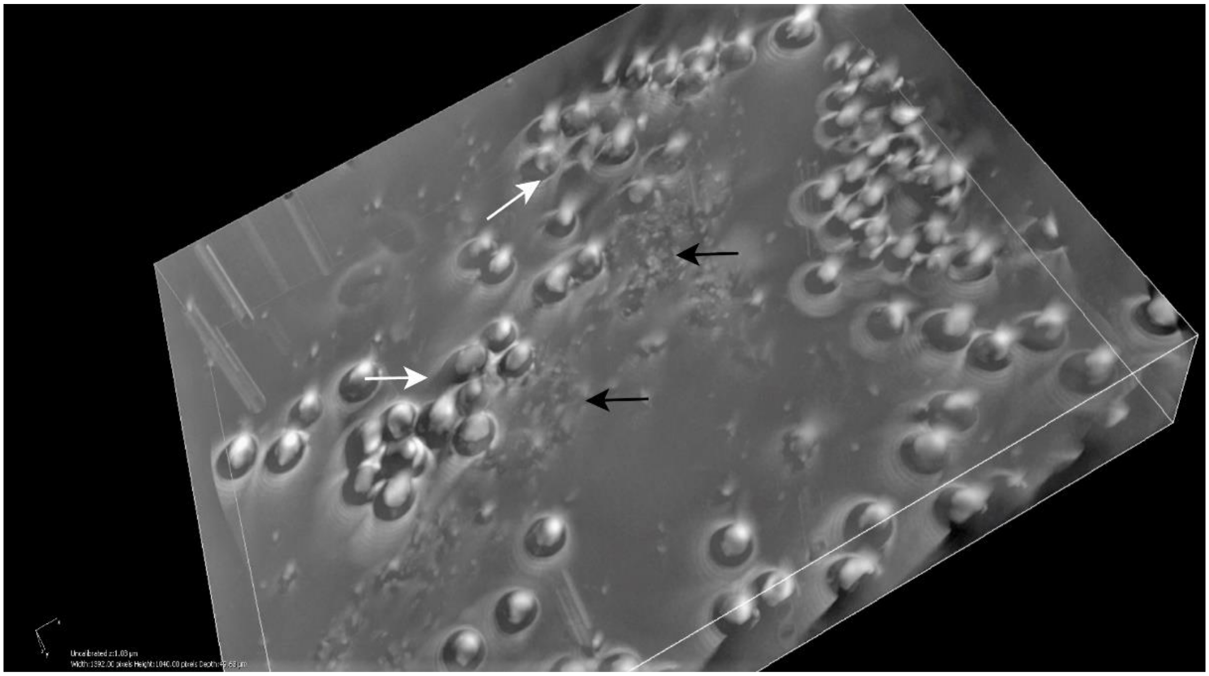

**Supplementary Fig. 3** 3D composite image of co-culture in hydrogel. White arrows show the *M. kaistiae* aggregates above the bacterial SIO_La6 aggregates (black arrows). (Brightfield magnification 200x)
